# Branching Networks Can Have Opposing Influences on Genetic Variation in Riverine Metapopulations

**DOI:** 10.1101/550194

**Authors:** Ming-Chih Chiu, Bin Li, Kei Nukazawa, Vincent H. Resh, Thaddeus Carvajal, Kozo Watanabe

## Abstract

**Aim:** Fractal networks, represented by branching complexity in rivers, are ubiquitous in nature. In rivers, the number of either distal (e.g., in headwater streams) or confluent (e.g., in mainstems) locations can be increased along with their branching complexity. Distal- or confluent-spatial locations can result in fewer or greater corridor linkages that can alter genetic divergence at the metapopulation scale. These mechanisms underlying the resulting genetic structuring remain poorly understood at the metapopulation scale, particularly in terms of the roles of species-specific dispersal traits. The objective of this study is to mechanistically understand how branching complexity can simultaneously influence genetic divergence in opposite directions.

**Location:** Northeastern Japan

**Methods:** To evaluate the integrated influences of network complexity and species dispersal on genetic divergence among populations at the catchment scale, we conducted simulation modelling on a mechanistic framework based on Bayesian inference by adapting empirical genetic data from four macroinvertebrate species. Simulations were then done using empirical and virtual species-characteristics on virtual river networks.

**Results:** Our novel simulation showed that both greater landscape connectivity (resulting from shorter watercourse distance) and greater isolation of distal locations occurred in the more-branched river networks. These two spatial features have negative and positive influences on genetic divergence, with their relative importance varying among different species and dispersal characteristics. Specifically, genetic divergence at the metapopulation scale increased for species having higher downstream-biased dispersal but decreased for species having higher upstream-biased dispersal. Distal populations (e.g., in headwaters) have higher genetic independence when downstream-biased asymmetry is higher.

**Main conclusions:** We found a strong association between species dispersal and evolutionary processes such as gene flow and genetic drift. This association mediates the pervasive influences of branching complexity on genetic-divergence in the metapopulation. It also highlights the importance of considering species dispersal-patterns when developing management strategies in the face of rapid environmental-change scenarios.

## Introduction

There is increasing interest in understanding how landscape architecture determines spatial patterns of intraspecific genetic-diversity (Paz-Vinas, Loot, Stevens, & Blanchet, 2015; Phillipsen & Lytle, 2013), although a mechanistic understanding of this process remains highly challenging in the real world. For example, ecological and evolutionary evidence and theories that are derived from simplified landscapes are insufficient for understanding spatial genetic patterns in complex systems, such as rivers (Fourcade, Chaput-Bardy, Secondi, Fleurant, & Lemaire, 2013; Thomaz, Christie, & Knowles, 2016). More in-depth explorations of the integrated genetic effects of species dispersal and landscape connectivity on metapopulations (here defined as groups of subpopulations with dispersal interactions) occurring in complex habitats are needed. These efforts are especially important when fragmented landscapes that result from global-change influences affect spatial genetic variability (Martins et al., 2016; Prunier, Dubut, Loot, Tudesque, & Blanchet, 2018).

In nature, fractal branching-networks (e.g., those with “treelike” patterns) have similar structural features to those seen in fluvial landscapes (Green, Klomp, Rimmington, & Sadedin, 2006). In these environments, landscape connectivity shapes evolutionary processes, such as gene flow and genetic drift, which ultimately produce the spatial patterns observed in intraspecific genetic variation (McRae, 2006; Phillipsen et al., 2015; Thomaz et al., 2016). In analyzing the evolutionary action of landscape architecture, branching networks can be characterised by being either distal or confluent locations (e.g., in headwater streams or mainstems, respectively), which allow fewer or greater corridor linkages, respectively. Therefore, these distal and confluent locations may have opposing influences by increasing or decreasing genetic divergence at the metapopulation scale, respectively (Alp, Keller, Westram, & Robinson, 2012).

The tendency of species to disperse among habitats is not necessarily occurring in a single direction (i.e., toward distal or confluent areas) or in different directions. These differences can result in dispersal asymmetry that can dictate the isolation processes between pairs of populations within a metapopulation (Junker et al., 2012; Ma, Shen, Bearup, Fagan, & Liao, 2020; Morrissey & de Kerckhove, 2009). For example, riverine populations occurring in a network’s distal branches (e.g., different headwaters of a riverine system) can still be connected to a common source population in downstream confluences through dispersal. However, downstream-biased dispersal (a tendency for higher dispersal downstream than upstream) may lead to weak connections among headwaters and result in a large genetic divergence among riverine species of fish and macroinvertebrates (Alp et al., 2012; Paz-Vinas & Blanchet, 2015; Paz-Vinas, Quéméré, Chikhi, Loot, & Blanchet, 2013). Theoretically, more isolated tributaries within a highly-branched network would result in higher genetic differentiation between local populations (Thomaz et al., 2016).

In contrast to the effects of dispersal asymmetry, network branching (e.g., in river systems) can enhance connectivity between populations by naturally increasing the number of confluences and by shortening the distance of their pathways (Labonne, Ravigné, Parisi, & Gaucherel, 2008; Paz-Vinas & Blanchet, 2015). Stream-dwelling species with strong tendencies to migrate upstream, such as aquatic insects that disperse by flying during their terrestrial adult stages (Petersen, Masters, Hildrew, & Ormerod, 2004; Winterbourn, Chadderton, Entrekin, Tank, & Harding, 2007), may exhibit low downstream- or upstream-biased gene flow. In this case, there is weaker isolation between distal populations in the river network when these populations receive more gene flow because of their shared source-population that occurs at downstream confluences (Alp et al., 2012). Therefore, dispersal asymmetry provides a mechanism that can underlie the widely observed patterns of spatial genetic diversity and genetic differentiation that is found in river systems (Kawecki & Holt, 2002; Pilger, Gido, Propst, Whitney, & Turner, 2017).

Based on simulation modelling, the objective of this study is to mechanistically understand how branching complexity can influence genetic divergence, and how it can do this in opposing ways depending on dispersal traits. Although dendritic structure and species dispersal have been reported to determine spatial genetic patterns across a river network in Switzerland (e.g., Alp et al., 2012), our study extends this observation to the metapopulation level through the comparison of multiple networks having varying levels of branching complexity (e.g., as in Thomaz et al., 2016). Through empirical data of riverine macroinvertebrates in northeastern Japan, we constructed the genetic simulations using empirical and virtual species-characteristics on virtual river networks. Herein, we hypothesised that increased network branching has positive effects on genetic divergence at the metapopulation scale for species with downstream-biased dispersal but has the opposite, negative effect for those with symmetric dispersal or upstream-biased asymmetric dispersal. This difference implies that the consequences of network branching, namely landscape connectivity (via watercourse distance) and the isolation of distal locations (e.g., headwaters), may have opposite effects on genetic structure and attributes of the metapopulation.

## Materials and methods

### Framework overview

The opposing influences of different dispersal modes on the genetic divergence within a metapopulation can be theoretically explained but these influences have not been demonstrated to occur along a gradient of river branching in multiple networks. To demonstrate these influences, we followed two successive steps. First, by using a mechanistic model based on evolutionary processes and asymmetric dispersal, we explored the spatial genetic variation among four macroinvertebrate species with flying adult-stages that occur in a river network of northeastern Japan. Second, we evaluated how branching complexity of random virtual networks can differentially affect the broader genetic differentiation throughout catchments in the context of different asymmetric gene-flow modes.

To achieve the first step, we used the empirical data (see the subsequent section “*Empirical catchment and genetic data*’) and developed a metapopulation genetic-model based on isolation by distance (i.e., lower gene flow with greater separation in terms of distance along the watercourse) and asymmetric dispersal (i.e., upstream- and downstream-biased movements) between local populations in the river network (see the subsequent section “*Development of metapopulation genetic model*’). Rather than using their actual locations, the watercourse distances between local populations were used directly as the input data in the model. Consequently, if the spatial connectivity among some local populations is high, then their genetic structure would tend to have consistent differences from that of their metapopulation (their genetic pool). In turn, this model was fitted to these empirical data with its parameters estimated by Bayesian inference (see the subsequent section ‘*Bayesian fitting process*’). To validate this inference process using the approach of Zurell et al. (2010), we performed this metapopulation genetic modelling with known values of parameters to confirm whether the Bayesian estimates matched these known values (Appendix S1). The Bayesian filtering process can provide the reliable estimation of model parameters when these known values were within the 95% confidence interval of the posterior distribution of estimates. However, the variance parameters (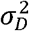 and 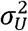) had relatively large uncertainty in the scenario of equal scale parameters (*c_D_ = c_U_*; Figure S2).

Bayesian estimation can provide realistic ranges of dispersal asymmetry and other parameters. In our model, these parameters were related to biological features of the four species and were not independent of each other. Because a particular species can be described as having a certain combination of parameters based on a set of trait characteristics and through its systematic clade, comparisons among these species can reveal the opposing influences of branching complexity on metapopulation genetic-structure. This can be done by examining the parameter space (i.e., the ranges of possible values of multiple parameters) and how each region of this parameter space reacts to different branching contexts. We used these ranges in our next step to achieve the main objective of this study, which was to evaluate the influence of network branching, and make our simulations be closer to the empirical system than to an arbitrary determination. The different parameter combinations (derived from Bayesian fitting process for the four macroinvertebrate species) allowed the creation of virtual species.

For the second step, the metapopulation genetic model constructed above was used to simulate the river branching influence on genetic divergence in different dispersal modes (i.e., by setting up the virtual species) in 1,000 virtual networks (see the subsequent section ‘*Simulation of river-branching influences*’). These virtual networks were randomly created to represent a variety of branching complexity.

### Empirical catchment and genetic data

In an empirical catchment in the Natori and Nanakita Rivers located in northeastern Japan (integrated catchment area c. 1200 km2; Figure 1), the flow regime exhibits a seasonal pattern, with spring flooding from snowmelt. In the integrated catchment, the rivers flow from the western headwaters at an elevation of 1350 m on Mount Kamuro to the east and into the Pacific Ocean. Approximately 60% of this area is forested and mountainous. Two major reservoir dams (Kamafusa and Okura dams) are located there on the rivers. The regional lowlands are farmlands (13%, primarily with rice paddy fields) and a mixture of residential and commercial areas (11%).

**Figure 1.**
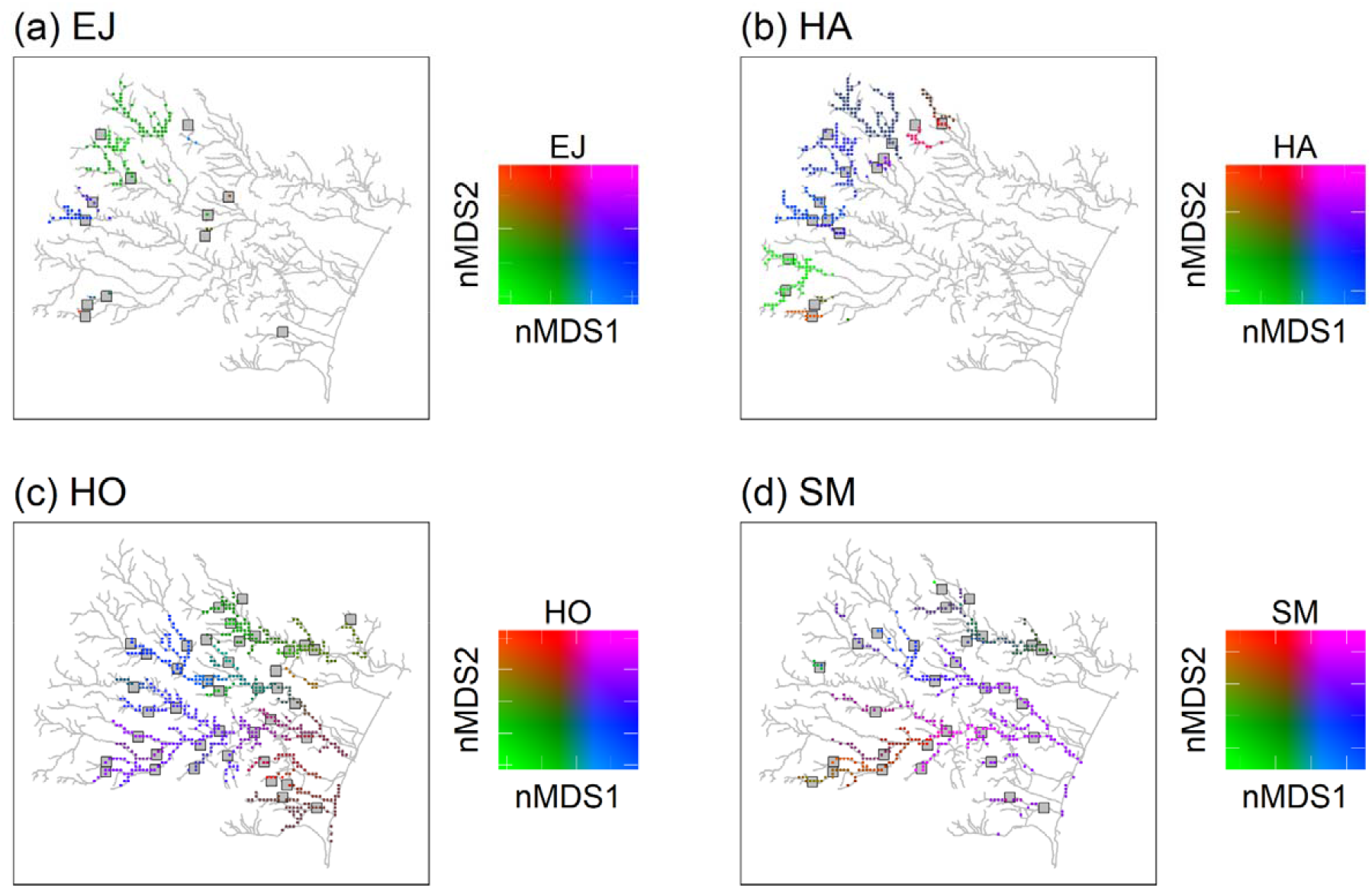
Study catchment area and distribution of (a) *Ephemera japonica* (EJ), (b) *Hydropsyche albicephala* (HA), (c) *Hydropsyche orientalis* (HO) and (d) *Stenopsyche marmorata* (SM), and the predicted genetic divergence of their local populations on the basis of nMDS (nonmetric multidimensional scaling) of the modeled pairwise *G_ST_* between river grid squares (500 × 500 m^2^) in northeastern Japan (see Appendix S1). Gray squares and color points indicate sampling sites and predictions, respectively.

Genetic data were generated from a previous study (Watanabe, Kazama, Omura, & Monaghan, 2014) We used neutral AFLP markers from four macroinvertebrate species (the mayfly *Ephemera japonica*, and the caddisflies *Hydropsyche orientalis, Stenopsyche marmorata*, and *Hydropsyche albicephala*) in this catchment (Appendix S1 and Figure S1). These species all have substantially diverse habitat specificity and distributions within the network (Nukazawa, Kazama, & Watanabe, 2015; Nukazawa, Kazama, & Watanabe, 2017; Watanabe et al., 2014).

### Development of metapopulation genetic model

This metapopulation genetic model has the two modules that correspond to observation and to evolutionary processes. The observation module describes the statistical inference of the true value of population’s genetic-frequency based on genetic measurements derived from our sampling of parts of individuals of population. We use Bayesian inference (see the subsequent section ‘*Bayesian fitting process*’) to fit the output of this module to our genetic measurements and then to calibrate the model parameters. By connecting statistical frameworks with two key processes of gene flow and genetic drift, the evolutionary module allows the genetic frequency of each local population to be equivalent to the frequency of the metapopulation with an estimate of the deviation of the frequency. The deviation is based on the genetic similarity of distance-decay among local populations, with the values of deviation tending to be similar if these populations have high ecological connectivity among them along watercourses. The paragraphs below describe the statistical inference used for the descriptions of these two modules.

The observation module describes how genetic measurements and sampling errors are produced. A single AFLP locus has two allelic types, labelled ‘1’ and ‘0’ for the dominant and recessive types, respectively. We denote by *z_k,l_* the number of individuals for which the dominant type ‘1’ corresponds to genotypes (‘1’, ‘1’) or (‘1’, ‘0’) in our diploid species, which was detected at locus *l* from the total number of individuals (*s_k_*; with types ‘1’ and ‘0’ together) in local population *k*. Here, *z_k,l_* was used as the modelling output calibrated by our observed data. This random sampling process can be characterised by a binomial distribution as follows (Guillot, Vitalis, Rouzic, & Gautier, 2014):

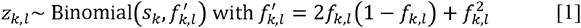

where, in this local population, the frequency of allele ‘1’ is denoted by *f_k,l_*. The allele frequencies are independent between loci. The detection probability 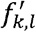 linked to *f_k,l_* was set up for the allele type ‘1’ corresponding to genotypes.

The evolutionary module describes the genetic structuring of local populations within a metapopulation. For each locus, the frequency of allele ‘1’ in the local populations is determined by their genetic variation (i.e., that related to genetic drift), the watercourse distance between local populations within the network (related to gene flow), and the allele frequencies of the metapopulation. Without genetic drift and natural selection, gene flow leads to genetic homogeneity among local populations, which results in allele frequencies at loci in local populations matching those of their metapopulation (Andrews, 2010). We denote the deviation from the expected frequency of allele ‘1’ in a logit scale as θ_*k,l*_, which was derived from the assumption of local populations being connected at the metapopulation scale. The allele frequency *f_k,l_* is obtained from the inverse logit transformation as follows:

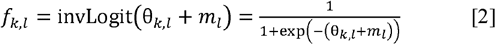

where *m_l_* denotes the metapopulation’s logit-transformed allele frequency. The deviation θ_*k,l*_ can be modelled by a multivariate normal distribution as follows (Bradburd, Ralph, & Coop, 2013):

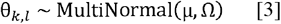

where μ denotes the vector of zero, and the covariance matrix Ω is a function of the watercourse distance between local populations and their spatial relationships. These spatial connections between local populations are either streamflow-connected (populations with downstream–upstream relationships) or streamflow-disconnected (e.g., populations in different headwaters). This statistical process allows similar values of deviation for local populations having high ecological connectivity along watercourses. We modelled the covariance across local populations as a function of the shortest watercourse distance, *h_ij_*, along the river network between populations *i* and *j* as follows (Ver Hoef & Peterson, 2010):

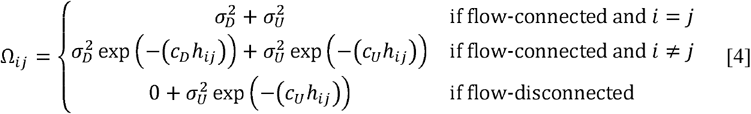

where the first or second addend (i.e., for any number that’s added to another; e.g., 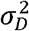 and 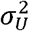 are different addends of their summation 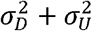) describes the autocovariance, with the variance 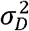 or 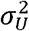 and the scale parameter *c_D_* or *c_U_* related to the downstream (D) or upstream (U) movement, respectively. In the first addend, the autocovariance is set to zero for any two streamflow-disconnected populations. In other words, streamflow-disconnected populations are assumed to be independent and have no gene flow between them via downstream movement. In the second addend, the autocovariance can apply to both situations. The genetic connections between streamflow-disconnected populations are attributed to the upstream gene-flow coming from a common source population in the downstream confluences. High values of variance and small values of the scale parameter lead to weak isolation as a function of the watercourse distance.

### Bayesian fitting process

In the Bayesian framework “Stan’ (Stan Development Team, 2018), the R interface “RStan’ (Stan Development Team, 2019) was used for genetic modelling of the metapopulation and to fit the model using our empirical data (see the above section ‘*Framework overview*’). For each species, four Markov Chain Monte Carlo (MCMC) chains (which were used for numerical approximations of Bayesian inference) were run with 20,000 iterations each, and the first half of the iterations for each chain were discarded as “burn-in”. This iteration setting was determined by the convergence of MCMC chains when the R-hat statistic of each parameter approached a value of 1. To estimate the model parameters, 2,000 samples obtained by collecting one sample out of every 20 iterations for each chain were used to build each parameter’s posterior distribution. The target average for the proposal acceptance-probability and initial discretization interval was set to 0.95 and 0.1, respectively, during Stan’s adaptation period.

Prior distributions were set up for the model parameters based on their data type (e.g., continuous or discrete), range of values (e.g., unlimited or positive), adjustments using an informative feature to keep reasonable values but not eliminating those that might be appropriate, parameter validation (through the virtual ecologist approach; see Appendix S1), and model-convergence efficacy. The transformed allele frequency *m_l_* has a normal distribution with mean of 0 and variance τ, and a half-Cauchy distributed hyper-prior on the variance τ with a location of 0 and a scale of 5. The variances (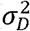 and 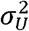) each have a half-Cauchy distribution with a location of 0 and a scale of 5. The scale parameters (*c_D_* and *c_U_*) each have a half-normal distribution with mean of 0 and variance of 2. The four macroinvertebrate species examined (*E. japonica, H. orientalis, S. marmorata* and *H. albicephala*) each had both types of relationship between locations (i.e., streamflow-connected and -disconnected pairs).

### Creation of virtual river networks

We created 1,000 virtual river networks with varying branching-complexities (Terui et al., 2018; Yeakel, Moore, Guimarães Jr., & de Aguiar, 2014). These virtual river networks (with scale length *e* equal to 1 km between local populations) have the branching probability *P* and metapopulation size *N* (which is an integer and represents the number of local populations in a river network) randomly drawn from 0 to 1 and from 100 to 500, respectively (Figure S3). This scale length was determined by referring to the local scale of stream macroinvertebrate occurrence, e.g., a 100 m reach (Stoll, Breyer, Tonkin, Früh, & Haase, 2016). Natural river networks in Japan have a range of *P* from 0.3 to 0.6 (Terui et al., 2018). The full range of *P* can reflect general conditions and provide insights into networks that, because of human intervention, depart from natural value of *P* (e.g., a network with *P* = 0.1 can represent a human-created canal; Figure S3).

The virtual river networks comprised nodes with scale length e, with each node representing a local population. These nodes were assigned to be either branching (or representing an upstream terminal) or non-branching with a probability of *P* or 1 – *P*, respectively. A segment was defined as a series of non-branching nodes terminating at a branching (or terminal) node. The number of nodes in a segment is a random variable that is assumed to be geometrically distributed with the probability of success *P* (equal to the branching probability). To achieve the target number of total nodes in a network, individual segments were formed by consecutively drawing their number of nodes from the geometric distribution. As a statistical reformulation of Horton’s laws, a geometric distribution has been used to describe the segment lengths (e.g., Peckham & Gupta, 1999). Before merging the segments to create a river network, the whole drawing process was repeated if a non-equal number of total nodes or a non-odd number of segments was attained. To create the river network, these segments were put together as a hierarchically merged pool in three steps (Figure S3). For Step 1, one segment, consisting of one or more nodes, was randomly selected as the root, and its upstream end was merged with the downstream end of two randomly selected segments. From this procedure, the semi-complete network had two unmerged upstream ends each for the next possible merger. For Step 2 Two more segments were then randomly selected, and their downstream ends were merged together to the random draw of one of two (or even more at subsequent steps) unmerged upstream ends of the semi-complete network. For Step 3, Step 2 was repeated until there were no available segments left in the pool.

### Simulation of river-branching influences

We conducted stochastic simulations to illustrate the genetic differentiation and diversity at the metapopulation scale under river branching. Our metapopulation genetic model for each species with the Bayesian-median estimates of the parameters added was then used to simulate allele frequencies at loci in local populations in each of the 1,000 random river networks (see the above section ‘*Creation of virtual river networks*’). The spatial genetic structure was summarized to the global *G_ST_* to be the metapopulation’s genetic divergence. In addition, the metapopulation’s genetic diversity was calculated as global *H_t_* (Nei, 1973). We performed this simulation using the R packages ‘stats’ and ‘base’ (R Core Team, 2018).

We then built a regression model based on gradient boosting (GB) (Appendix S1) for each of the four macroinvertebrate species, which identified the importance of the following spatial features: (1) the fraction of any two local populations being streamflow-disconnected in all possible combinations; (2) the mean watercourse distance between local populations under different levels of river branching; and (3) the metapopulation size (i.e., number of local populations) for genetic divergence (global *G_ST_*). These features were considered to be summarised aspects of complex spatial configuration of habitat network.

Through further simulations on the same set of river networks, we illustrated how the downstream and upstream dispersal-related parameters of the scales (*c_D_* and *c_U_*, respectively) influence river-branching effects on the genetic divergence of the metapopulation (global *G_ST_*) and genetic diversity (global *H_t_*). We considered 3 × 3 (nine total) combinations of the scale parameters, with each having three values, which were the maximum, median, and minimum values of range of their Bayesian pooled estimates where the posterior median of both *c_D_* and *c_U_* across the four macroinvertebrate species were pooled together. Because of the shared pool, the maximum, median, or minimum values set for the two scale parameters in the combinations are the same. We set the metapopulation’s logit-transformed frequency of allele “1” at each locus to zero during each simulation. The number of loci was set as the median across the four macroinvertebrate species. In addition, we set the variance parameter (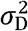 or 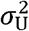) to the median value of their Bayesian pooled estimates, with the two variance parameters having the same values. We replicated the nine combinations in each of the 1,000 random river networks.

## Results

### Metapopulation genetic modelling

When our Bayesian model was fitted to the empirical genetic data in the Natori and Nanakita catchments, the *R*^2^ values derived from the residuals (i.e., the differences between the observed and predicted numbers of type ‘1’ alleles at a locus from the number of alleles observed in a local population; see Equation 1) were 0.98, 0.97, 0.97, and 0.93 for *H. orientalis, S. marmorata, H. albicephala* and *E. japonica*, respectively (Figure S4).

The metapopulation allele frequencies at loci are species-specific, and the variation of allele frequency was greater in the widespread *H. orientalis* and *S. marmorata* than in the other two species that had narrower habitat distributions (Figure S1). Pairwise genetic differences between empirical local populations tended to increase with their watercourse distances throughout the four macroinvertebrate species (Figure 1). Despite substantial variation in the scale parameter that amplified the isolating effect of distance across study species (Figure 2), there was a consistent decline in the genetic correlation between populations (the covariance Δ_*ij*_ divided by the variance 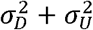 see Equation 4) with the increasing distance between local populations being obvious for the three caddisfly species but not the mayfly having lower value of *c_D_* (Figure S5).

**Figure 2.**
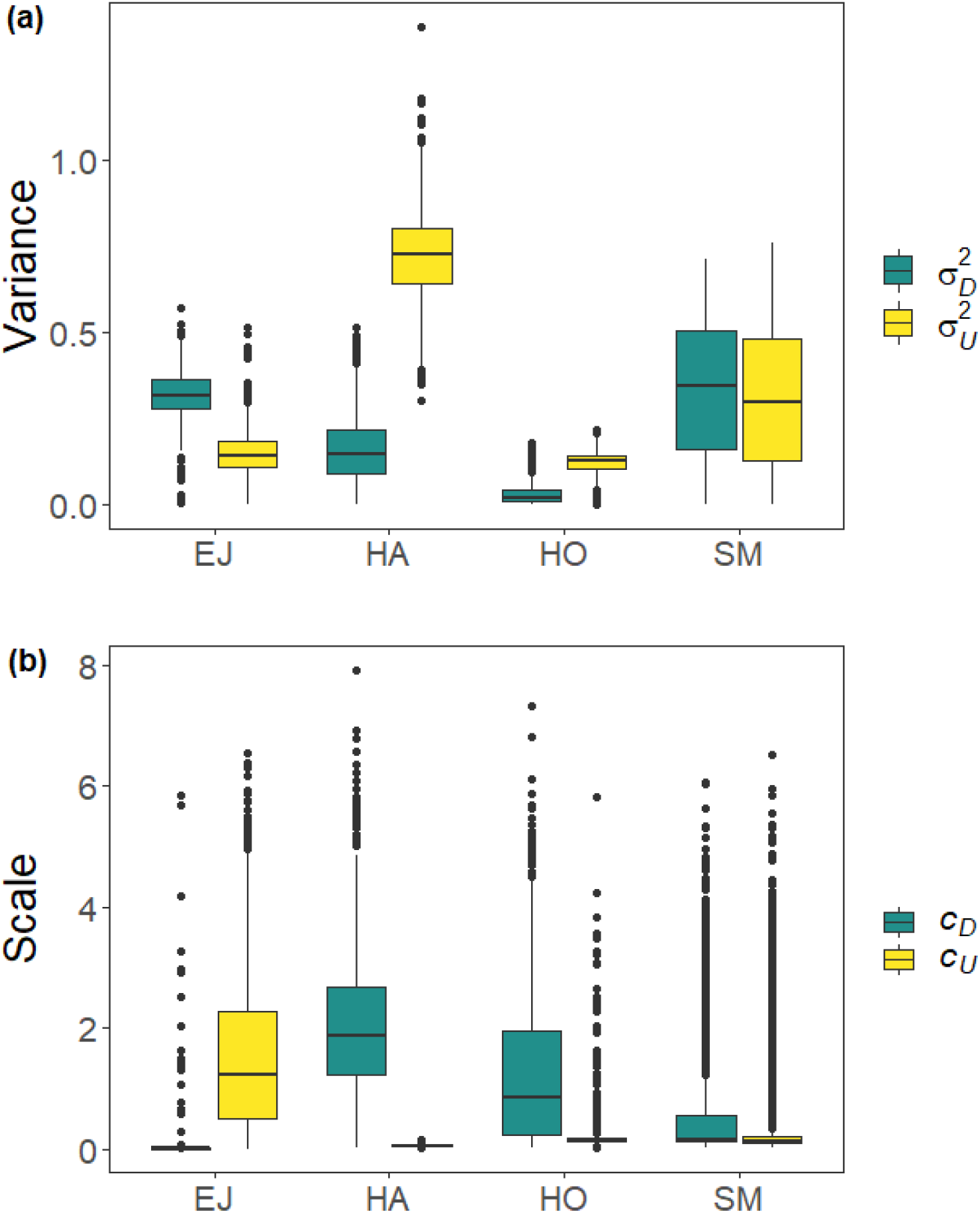
Estimated values of (a) variances (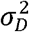 and 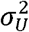) and (b) scales (*c_D_* and *c_U_*) of covariance function (see Equation 4) with the posterior distribution in our Bayesian modeling for *Ephemera japonica* (EJ), *Hydropsyche albicephala* (HA), *Hydropsyche orientalis* (HO), and *Stenopsyche marmorata* (SM).

### Simulation of river-branching influences

The increasing branching probability of river networks caused changes in the observed spatial features (Figure 3). For example, the fraction of any two local populations being streamflow-disconnected (e.g., occurring in different tributaries) across the metapopulations occurs at high rates in the heavily branched river networks. However, we found shorter watercourse distances occurred between local populations in river networks with higher branching probability. As the size of the virtual network increases, the metapopulation size (i.e., the total number of local populations of the network) increases as do the values of both spatial configurations under the same level of river branching.

**Figure 3.**
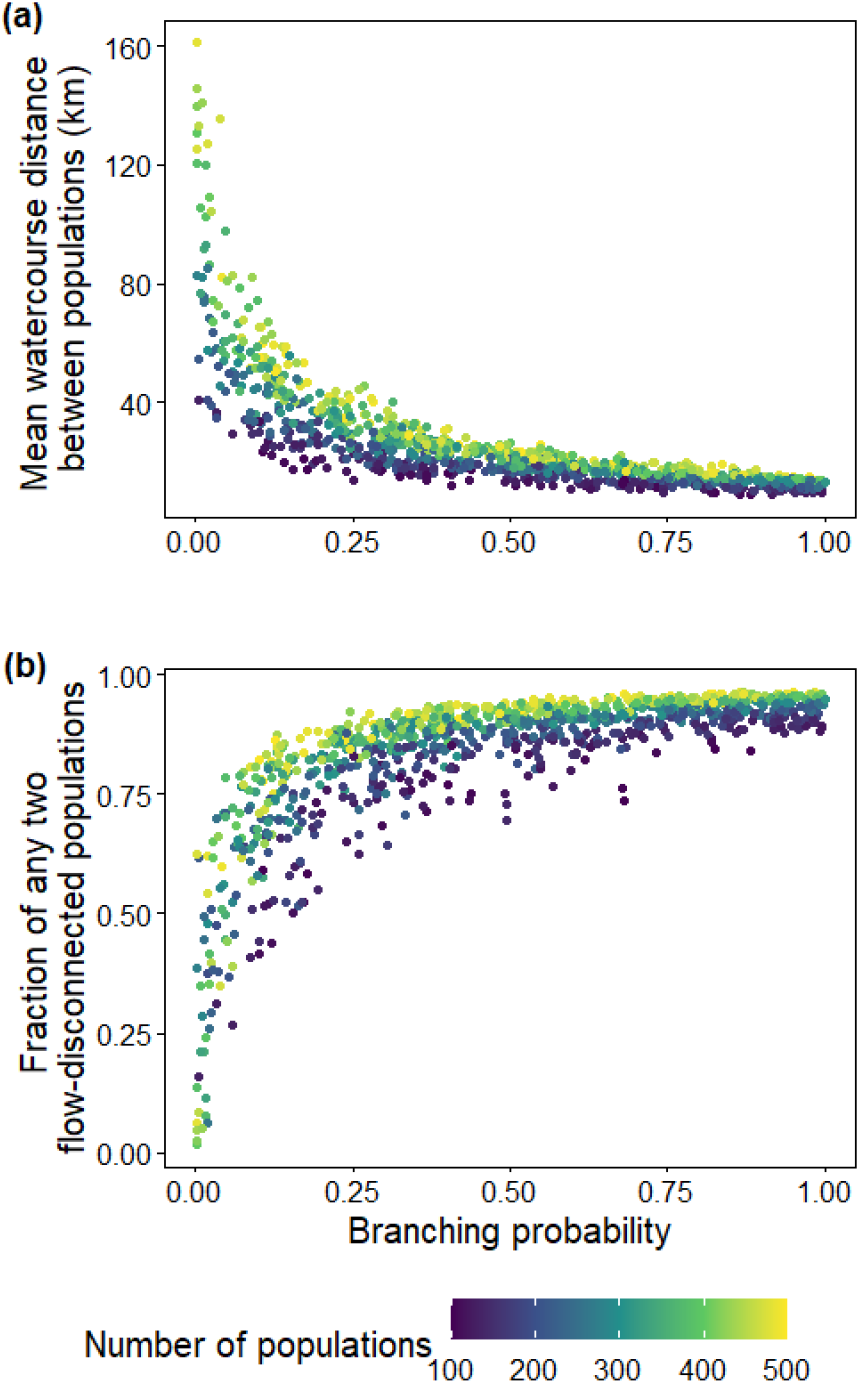
Theoretical predictions for relationships of (a) mean watercourse distance between populations or (b) fraction of any two streamflow-disconnected populations (e.g., in headwaters) in all combinations having branching complexity under differential metapopulation sizes (range: 100 to 500 local populations).

Two spatial features (watercourse distance and fraction of flow-disconnected sites; Figure 3) summarized the spatial configuration of habitat network and were linked to the genetic differentiation of metapopulations (*G_ST_*; Figure 4) and total genetic diversity (*H_t_*; Figure S6). However, species-specific responses to the influence of river branching were evident. For example, increased branching probability decreased the genetic divergence and diversity of the metapopulation for one caddisfly (*H. albicephala*), but the mayfly *E. japonica* showed the opposite response (higher genetic divergence) before a branching threshold. According to the GB modelling results, the relative importance of streamflow-disconnected habitats, compared to the landscape connectivity via a shorter watercourse distance, was higher for the mayfly than in all the caddisflies examined (Table 1).

**Table 1.** Performance measures and relative importance of predictors in gradient-boosting models for *Ephemera japonica* (EJ), *Stenopsyche marmorata* (SM), *Hydropsyche orientalis* (HO), and *Hydropsyche albicephala* (HA). The simulated genetic divergence of the metapopulation (global *G_ST_*) is the response variable and fraction for any two streamflow-disconnected local populations, the mean watercourse distance between populations, and metapopulation size, all of which are predictor variables.

| Species | Number of trees | Model performance |  | Relative importance (%) |  |  |
| --- | --- | --- | --- | --- | --- | --- |
|  |  | RMSE | R<sup>2</sup> | Streamflow-disconnected fraction | Watercourse distance | Metapopulation size |
| EJ | 3800 | 0.03 | 0.95 | 84.2 | 9.1 | 6.7 |
| HA | 950 | 0.05 | 0.89 | 1.4 | 95.9 | 2.7 |
| HO | 600 | 0.05 | 0.59 | 4.0 | 82.6 | 13.5 |
| SM | 600 | 0.04 | 0.50 | 6.6 | 26.0 | 67.4 |

**Figure 4.**
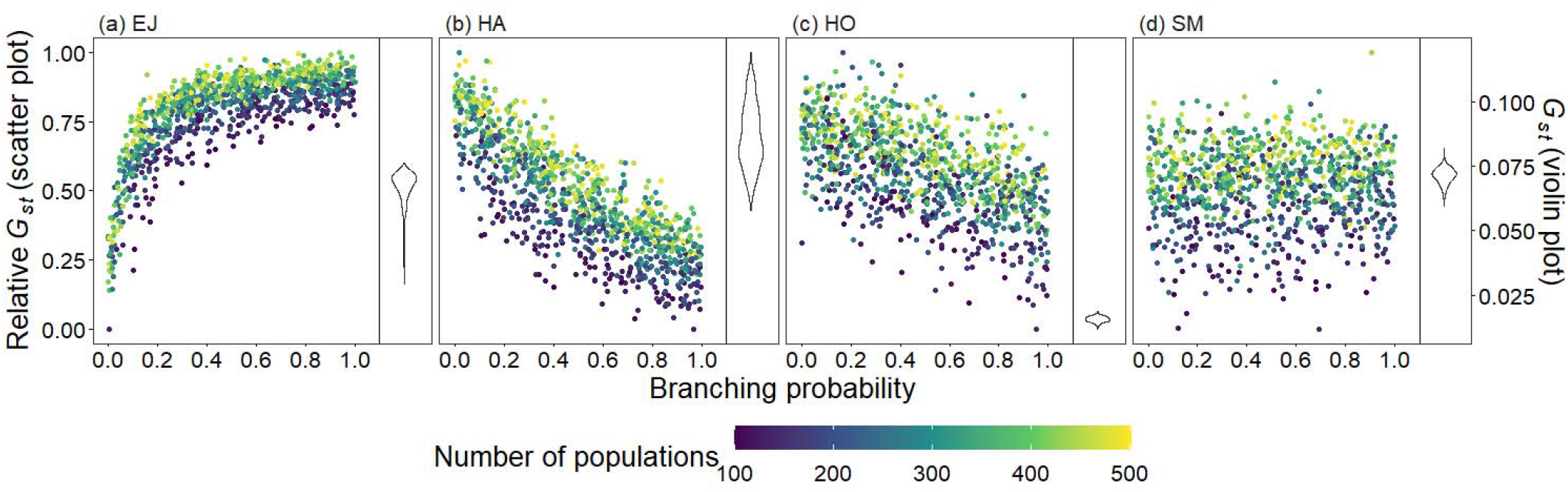
Theoretical predictions for relationships between metapopulation genetic divergence (global *G_ST_ = GG_ST_*) and branching complexity under different metapopulation sizes (range: 100 to 500 local populations) for (a) *Ephemera japonica* (EJ), (b) *Hydropsyche albicephala* (HA), (c) *Hydropsyche orientalis* (HO), and (d) *Stenopsyche marmorata* (SM). Relative *G G_ST_* = (*GG_ST_* - minimum *GG_ST_*) / (maximum *GG_ST_* - minimum *GG_ST_*) across the range for each species.

Branching complexity has various impacts on genetic divergence and diversity, which is determined by the relative values of the upstream and downstream scale-parameters in a genetic covariation function (Equation 4). Stronger dispersal tendency can be associated with a lower value of the scale parameter in either direction, which will result in weak isolation by watercourse distance within a network. When species dispersal was upstream-biased (*c_U_ < c_D_*), branching complexity enhanced genetic homogenization among local populations within a metapopulation. In addition, lower genetic divergence and diversity are related to lower downstream-biased and higher upstream-biased dispersals (relative higher *c_D_* and lower *c_U_*, respectively; Figures 5 and S7).

**Figure 5.**
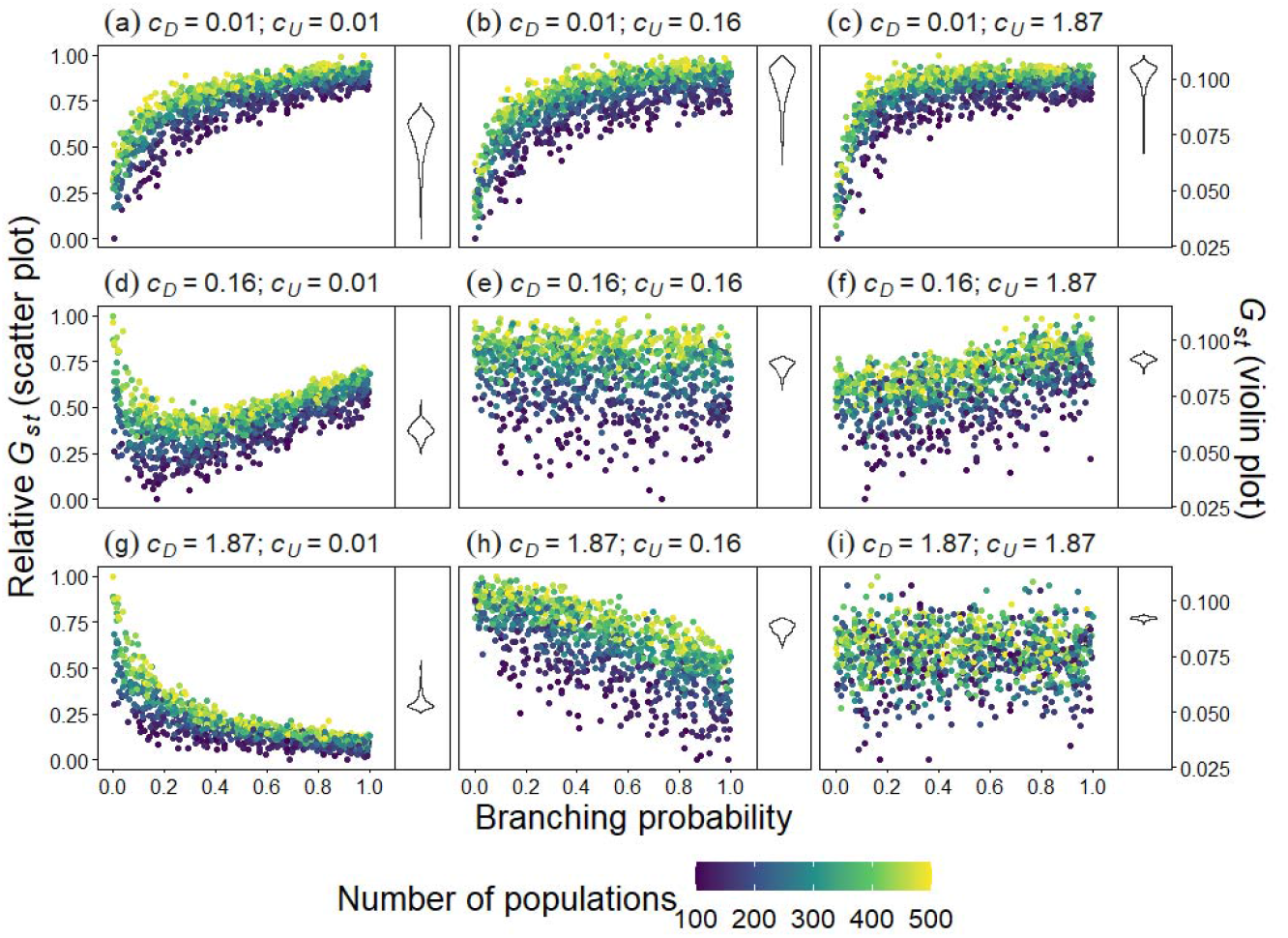
Theoretical predictions of relationships between metapopulation genetic divergence (global *G_ST_ = GG_ST_*) and branching complexity under differential metapopulation sizes (range: 100 to 500 local populations) for combinations of dispersal-related scale parameters in a genetic covariation function (Equation 4), including (a, e and i) *c_D_ = c_U_*, (b, c and f) *c_D_ < c_U_*, and (d, g and h) *c_D_ > c_U_*. The three values of each parameter are the maximum, median, and minimum values of range of Bayesian pooled estimates when the posterior median of both *c_D_* and *c_U_* across the four macroinvertebrate species were pooled. Relative *GG_ST_* = (*GG_ST_* - minimum *GG_ST_*) / (maximum *GG_ST_* - minimum *GG_ST_*) across the range for each parameter combination.

## Discussion

Using simulation modelling in virtual networks, we explored the integrated role of landscape architecture and species dispersal asymmetry in shaping genetic divergence at the metapopulation scale. We compared the spatial patterns of genetic differentiation of four macroinvertebrate species in river networks based on our Bayesian model that explicitly accounted for the effects of evolutionary processes among components of the metapopulations on their spatial genetic-structure. Our simulations indicated that greater landscape connectivity (via shortened watercourse distance) and higher distal isolation (e.g., in headwater streams) simultaneously occur in more-branched river networks and have countervailing influences on genetic divergence.

The two simplified, correlated aspects of complex networks (Figure S8) can have nonequal importance for different species and this difference can be linked to varying genetic divergence in the metapopulation. These results, derived from the simulation of virtual networks, may not directly apply to all empirical systems but they can provide hypothesized expectations for the relationship between genetic divergence and network branching. We suggest that species dispersal can cause varying levels of overall genetic differentiation in river networks having increased river branching, and can have either different or even opposite effects. We believe that this result can provide insights into other complex networks (e.g., highly fragmented landscapes or those with corridors via ocean and atmospheric circulation). In addition, these results highlight the fundamental importance of considering dispersal traits of individual species, which can result in different contributions to genetic connectivity, for the successful management of ecological corridors. Our theoretical evidence suggests the value of conducting further studies using empirical comparisons across multiple natural networks based on hypotheses generated from these modelling efforts.

### Branching complexity and genetic divergence

The role of network branching has been documented, to some extent, in genetic studies of riverscape conducted in situations where higher genetic diversity is observed in downstream populations than in upstream ones (Paz-Vinas et al., 2015). Also, greater river branching can increase the differences between such populations (Thomaz et al., 2016). In dendritic networks, our simulation results showed a species-dependent change in global genetic-differentiation levels that occur with increases in network complexity. This finding was also previously observed for two sympatric salmonid species that were found to have remarkably different spawning locations, mating systems, and population sizes, and for which biological traits mediated the influences of riverscape features that shape their dispersal and genetic divergence (Whiteley, Spruell, & Allendorf, 2004).

Little or no gene flow can occur with high network branching. This pattern of gene flow is caused by strong genetic isolation from an increase in the number of distal populations (e.g., in headwaters) when the same number of nodes in a habitat network is present. This pattern can generally be observed for some riverine species with high or intermediate levels of downstream-biased vagility, such as fish and molluscs (Osborne, Perkin, Gido, & Turner, 2014; Pilger et al., 2017; Terui et al., 2014). In one, early theoretical study that did not consider dispersal asymmetry (which would be analogous to having equal downstream and upstream dispersals in our study), the dendritic network structure was also documented to promote low genetic distances under conditions of high connectivity (Labonne et al., 2008). However, our results show that the different downstream and upstream gene flows considered in our model can act together to generate similar or even different relationships.

### Importance of species dispersal

The architecture of river networks can be an important extrinsic factor in explaining the observed and simulated genetic patterns in our study, but there was high variation among the species with different dispersal traits in our study. By adopting our mechanistic model that was fitted by empirical data on macroinvertebrate species with flying adult-stages, our simulations reveal that their better dispersal abilities can overcome catchment constraints, leading to low downstream-biased asymmetry. The opposite (negative) influence occurs under increased river branching. In addition, our comprehensive consideration of various branching patterns in river-network topologies in our simulations demonstrate the existence of opposing influences that can co-occur under branching complexity.

In the river systems we examined, asymmetric (either downstream- or upstream-biased) dispersal could determine the effect of branching complexity on genetic divergence of a metapopulation. In our model, the scale parameters for upstream- and downstream-dispersal tendencies (in the genetic covariation function; Equation 4) can be linked to dispersal asymmetry. These two factors together provide a mechanism behind the countervailing influences of river branching on the genetic divergence of metapopulations. Stronger dispersal can be associated with a lower value of the scale parameter either in downstream or upstream direction, which will result in weak isolation by distance within a metapopulation.

Our results suggested that, in the mayfly species examined (*E. japonica*) having high downstream-biased gene flow (based on a higher value of the scale parameter for upstream than for downstream dispersal), river branching has a positive influence on genetic divergence. This phenomenon can be attributed to a higher number of isolated distal branches in river networks (e.g., more headwaters) occurring under this dispersal asymmetry, as typically shown in fish (e.g., Pilger et al., 2017). Larvae of mayflies also are susceptible to high levels of drift during times of increased river flow and therefore have high potential to be strong downstream dispersers (Nukazawa et al., 2017).

In our system, an opposite (negative) influence of the branching network on the three caddisfly species with dispersal symmetry or even upstream-biased dispersal (based on the scale parameter for upstream being similar to or higher than that for downstream dispersal, respectively; Figures 2 and 4) was evident. This result could be attributable to their flying adults generally providing a wide dispersal range and strong terrestrial movement (e.g., at least in the upstream direction, which would compensate for downstream drift dispersal). This upstream movement dampened the isolation between distal branches in river networks for caddisflies compared to mayflies (and although not examined, likely stoneflies as well), which exhibit restricted distributions to areas very close to their sources of emergence in a stream (Winterbourn et al., 2007).

### Unexplained genetic variations

The Bayesian fitting process is more likely to have failed in retrieving the assumed value of variance parameters in the scenario of equal values of scale parameters than in other scenarios. These stochastic effects are derived from evolutionary processes modelled by autocovariances related to upstream and downstream dispersal along the habitat networks. The deviations from stochasticity are not significant when compared to the large difference between the actions of the autocovariances on the genetic structures. Thus, the Bayesian estimations are more easily confounded when both autocovariances have equal or similar influence on the stochastic processes. In general, more sampling of sites in the river network and additional collections of individuals from local populations would result in higher constraints for parameter estimations, which can decrease the stochastic effects. In river systems, this issue can occur from aquatic insects having different aerial-dispersal abilities and may not apply in other riverine species (e.g., fish and mollusc species) having downstream-biased dispersal.

Some bias in our modelling results could result from not considering overland dispersal, particularly when asymmetric dispersals along the watercourse in our model explain a large part of the genetic variation at a metapopulation scale. Overland dispersal across tributaries or catchments has been demonstrated to influence genetic differentiation (Alp et al., 2012; Chaput-Bardy, Fleurant, Lemaire, & Secondi, 2009). However, this type of dispersal may not have made an important contribution in our empirical system when compared with in-stream dispersal. Overland movement can be particularly important to strong fliers like damselflies and riverine birds, e.g., dippers (Hernández et al., 2012). However, aquatic insects with weaker flying strength (e.g., mayflies or caddisflies) have larger longitudinal tendency to use stream corridors than to use lateral areas away from the stream channel (Petersen et al., 2004). However, empirical genetic-evidence indicates that in-stream dispersal was the main dispersal pathway (Chaput-Bardy, Lemaire, Picard, & Secondi, 2008).

### Overview and management implications

From a conservation and management perspective that attempts to include considerations of the spatial genetic-structure of metapopulations (e.g., Luque, Saura, & Fortin, 2012), an understanding of the branching structure’s role in driving metapopulation genetic divergence is critical. Dispersal can dictate differences in genetic diversification in landscapes (Medina, Cooke, & Ord, 2018) and our predictive modelling based on asymmetric dispersals across networks sheds light on better management practices. For example, in the context of managing native or even invasive species (Chaput-Bardy, Alcala, Secondi, & Vuilleumier, 2017), our simulation results provide insight into the evolutionary importance of dispersal abilities and modes, suggesting that these intrinsic factors should be considered in decision-making processes where one management strategy does not fit all species (Figure S9). For example, the same management and conservation practices can produce different or even opposite results for species with varying levels of asymmetric gene flow and genetic drift (e.g., in dendritic river systems). Moreover, by adopting a Bayesian fitted mechanistic framework into local landscapes, simulations in virtual networks could result in insights gained via experimental or empirical studies that can assist managers build specific strategies for different regions. For example, the potential genetic influences from past or planned human alterations of river systems (e.g., dam construction, loss of headwater habitats from pollution, branches diverted for agriculture, or creation of flood-bypass channels) may be incorporated into assessments of branching complexity. In doing the assessments, ecological connections rather than purely hydrological linkages are considered to be the elements forming the structure of habitat networks. In addition, this approach can be considered for incorporation into management strategies (e.g., in-stream barrier removal and habitat restoration placed in the context of the river network) and anticipated responses to environmental changes (e.g., drying out of segments due to reduced precipitation under climate change).

## Data Accessibility Statement

No new genetic data were created. AFLP genetic data, including neutral and non-neutral loci, are available from the Dryad Digital Repository at: http://dx.doi.org/10.5061/dryad.82br3. Stan code is uploaded as a separate supplemental file.

## Acknowledgements

We are grateful to Maribet Gamboa for her constructive comments on the manuscript. This study was partly funded by the Japan Society for the Promotion of Science (JSPS). MCC was supported by the JSPS (JSPS Invitation Fellowship Program; L18522) and the Chinese Academy of Sciences (CAS Taiwan Young Talent Programme; 2017TW2SA0004).

## Author Contributions

All authors conceived the research. MCC, BI, KN, and TC performed analyses, and MCC performed modeling work. MCC, KW, and VHR wrote the draft of the manuscript, and all authors contributed substantially to revisions.

