## Appendix S1 for "Branching Networks Can Have Opposing Influences on Genetic Variation in Riverine Metapopulations"

**Extended Methods**

*Genetic data*

Genetic data were generated from a previous study (Watanabe *et al.*, 2014), and we briefly describe their collection here. For both empirical and theoretical analyses, we used genetic data of neutral amplified fragment length polymorphism (AFLP) markers from four macroinvertebrate species in this catchment. Three species were caddisflies (Insecta: Trichoptera), namely, *Hydropsyche orientalis*, *Stenopsyche marmorata*, and *Hydropsyche albicephala*, while the fourth was a mayfly (Insecta: Ephemeroptera), *Ephemera japonica*. In this integrated catchment, the species distributions varied considerably, from the widespread *H*. *orientalis* and *S. marmorata* to the narrowly distributed *E*. *japonica*, and *H. albicephala* (Figure 1). These species have similar ecological functions in river ecosystems by feeding on organic matter. Approximately 18 to 20 individuals collected at each sampling site were genotyped (12, 15, 41, and 30 sampling sites for *E*. *japonica*, *H*. *albicephala*, *H*. *orientalis* and *S*. *marmorata*, respectively). There were 473, 128, 129, and 220 polymorphic AFLP loci for *E*. *japonica*, *H*. *albicephala*, *H*. *orientalis* and *S*. *marmorata*, respectively. Based on the locus-specific genetic differentiation across this catchment, non-neutral loci identified by the software DFDIST (Beaumont & Nichols, 1996) and/or BayeScan (Foll & Gaggiotti, 2008) were removed, and 449, 111, 98, and 162 neutral AFLP loci for *E*. *japonica*, *H*. *albicephala*, *H*. *orientalis*, and *S*. *marmorata*, respectively, were retained and used for this study (Figure S1). Detailed protocols on the identification of non-neutral loci are described by Watanabe *et al.* (2014).

*Genetic prediction in this Japanese watershed*

We predicted genetic divergence of local populations across this Japanese watershed for *E*. *japonica*, *H*. *albicephala*, *H*. *orientalis*, and *S*. *marmorata* based on non-metric multidimensional scaling (nMDS) of the pairwise $G_{ST}$ between river grid squares (500 m × 500 m) (see Figure 1). The pairwise $G_{ST}$ (Nei, 1972) between all of the grid squares was calculated using “phylip” (Felsenstein, 1989) and its R interface “Rphylip” (Revell & Chamberlain, 2014). Based on covariance function with Bayesian estimation of parameters (see Figure 2), we performed the spatial prediction by Kriging interpolation using the R package ”gstat“ (Pebesma, 2004; Pebesma & Heuvelink, 2016). We removed part of river grid squares in prediction when they had no genetic correlations greater than 0.3 at any of the observed locations (based on our genetic modeling) and/or were classified as non-habitat (based on ecological niche modeling). An ecological niche modeling, based on habitat distribution (i.e., spatial allocation of species presence/absence) along environmental gradients, was performed using a “random forest” approach with the R package “randomForest” (Breiman, 2001) for determining habitat suitability in the watershed (Nukazawa K. et al., unpublished work). In brief, the model consisted of hydraulic and thermal predictors (e.g., current velocity and water temperature) computed using a hydrothermal simulation (Nukazawa, Kazama & Watanabe, 2015), and performance of the model’s classification for the four taxa ranged from excellent to fair based on independent validations with area under the curve (Swets, 1988). We used the R package “vegan” (Oksanen *et al.*, 2016) to perform nMDS.

*Gradient boosting (GB)*

GBs are a type of machine-learning algorithms used for analysing unilinear relationships at the base of multiple decision trees (Friedman, 2001). A decision tree is a non-parametric model used for classification or regression. In the boosting process, each of the next tree-models generated is added to improve on the performance of the previous ensemble of models by minimising deviance. Our GB modelling was performed using the R package ‘gbm’ (Greenwell *et al.*, 2018), in which the genetic divergence and other factors (the river features and metapopulation size) were dependent and independent variables, respectively. We used the R package ‘dismo’ to assess the optimal number of boosting trees via a cross-validation procedure (Hijmans *et al.*, 2017).

*Validation of fitted model parameters*

We performed a virtual ecologist approach (Zurell *et al.*, 2010) to confirm the validation of the posterior distributions of model parameters. First, we simulated particular scenarios as virtual truths based on our metapopulation genetic model. We considered 3 combinations of the scale parameters ( $c_{D}=c_{U}$, $c_{D}<c_{U}$, and $c_{D}>c_{U}$), each in a random network among 1,000 virtual river networks (see the section ‘*Creation of* *virtual river networks*’). The different parameter combinations characterized the virtual species for the virtual ecologist approach. In addition, 40 nodes were randomly selected as local populations across each network. The values of $c_{D}$ and $c_{U}$ in the combinations ( $c_{D}=c_{U}$, $c_{D}<c_{U}$, or $c_{D}>c_{U}$) were set to 0.25 and 0.25, 0.25 and 2.5, or 2.5 and 0.25, respectively. For each combination, we set the metapopulation’s logit-transformed frequency of the allele “1” at each locus to zero. In addition, the number of loci and variance parameter ($\sigma_{D}^{2}$ or $\sigma_{U}^{2}$) were set to 150 and 0.1, respectively.

Second, we sampled the emerging results based on our modelled observation-process (see Equation 4), and then fitted the models to this sampled data to confirm whether the Bayesian process could detect the set values of the model parameters in the different scenarios. For each scenario, four MCMC chains were run with 10,000 iterations for each to achieve the convergence (when the R-hat statistic of each parameter approached a value of 1). The first half of the iterations for each chain were discarded as “burn-in”, and 2,000 samples (obtained by collecting one sample every 10 iterations for each chain) were used to build each parameter’s posterior distribution. The target average for the proposal acceptance-probability and initial discretization interval was set to 0.95 and 0.1, respectively, during Stan's adaptation period.

**Supplemental Figures and Tables**

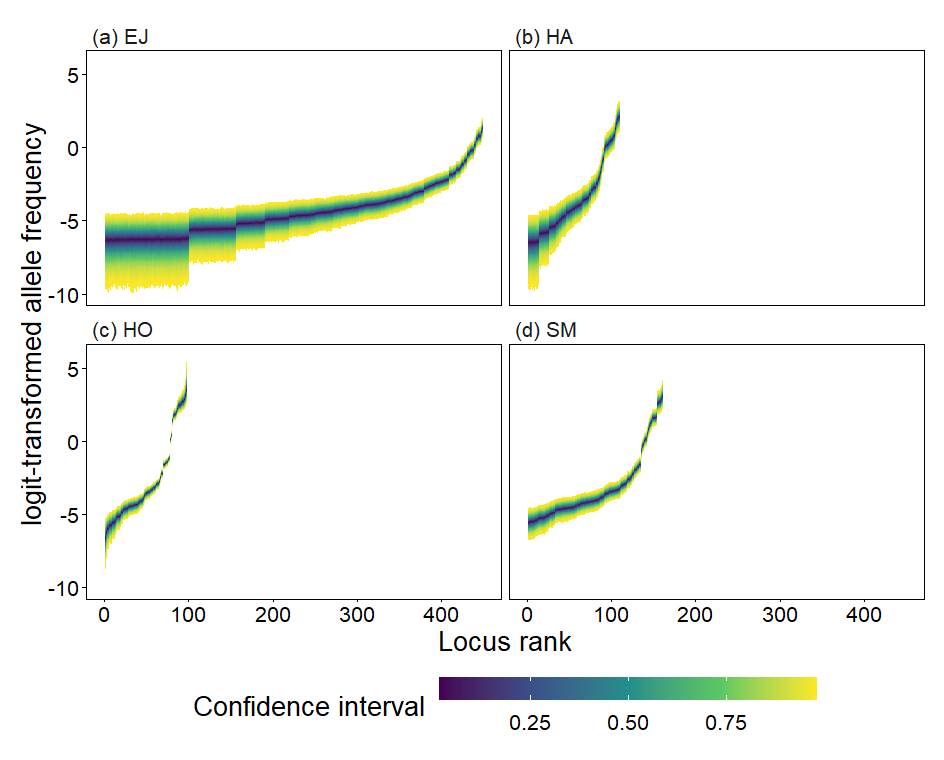

**Figure S1.** Metapopulation’s logit-transformed allele frequencies at loci with the posterior distribution in Bayesian modeling for (a) *Ephemera japonica* (EJ), (b) *Hydropsyche albicephala* (HA), (c) *Hydropsyche orientalis* (HO), and (d) *Stenopsyche marmorata* (SM). The range between upper and lower ends with the same color indicate one level of a confidence interval of the posterior distribution. For example, the internal line with a deep blue color shows the median (0% confidence interval).

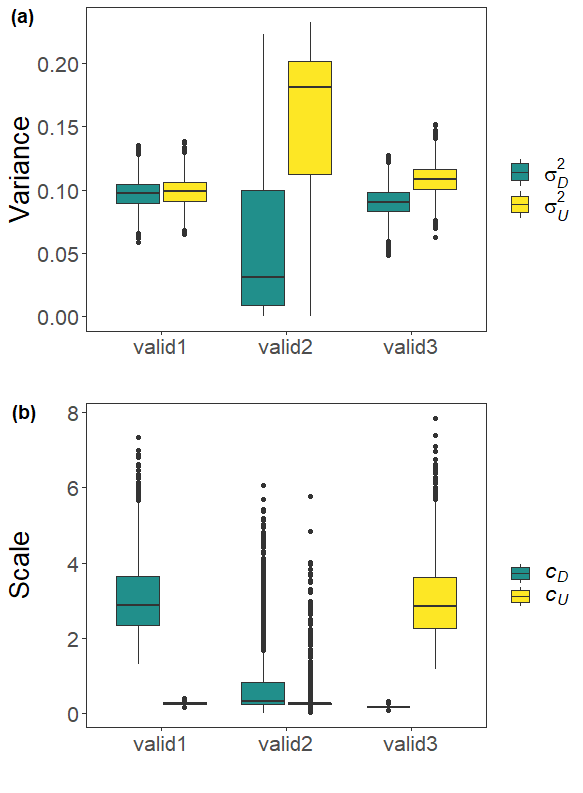

**Figure S2.** Estimated values of (a) variances ($\sigma_{D}^{2}$ and $\sigma_{U}^{2}$) and (b) scales ($c_{D}$ and $c_{U}$) of covariance function (see Equation 4) with the posterior distribution in our Bayesian modeling for scenarios of the virtual ecologist approach (valid1: $c_{D}>c_{U}$ ; valid2: $c_{D}=c_{U}$, valid3: $c_{D}<c_{U}$).

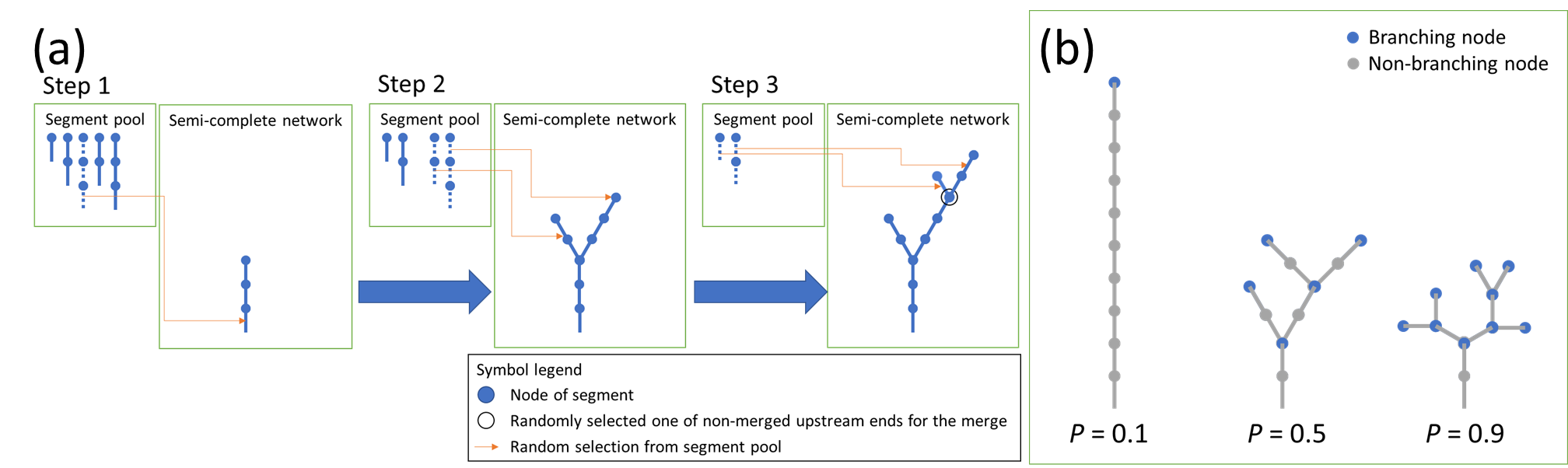

**Figure S3.** (a) Synthesis of the generation process of a river network based on an example pool of five segments, each with a series of non-branching nodes terminated at a branching (or terminal) node. These segments are selected subsequently and then merged hierarchically; (b) Examples of random river networks with ten nodes each assigned to be either a branching (or representing an upstream terminal) one with probability *P* or a nonbranching one with probability 1 − *P*. The virtual river networks comprised nodes with scale length (e.g., equal to 1 km between two adjacent nodes), with each node representing a local population.

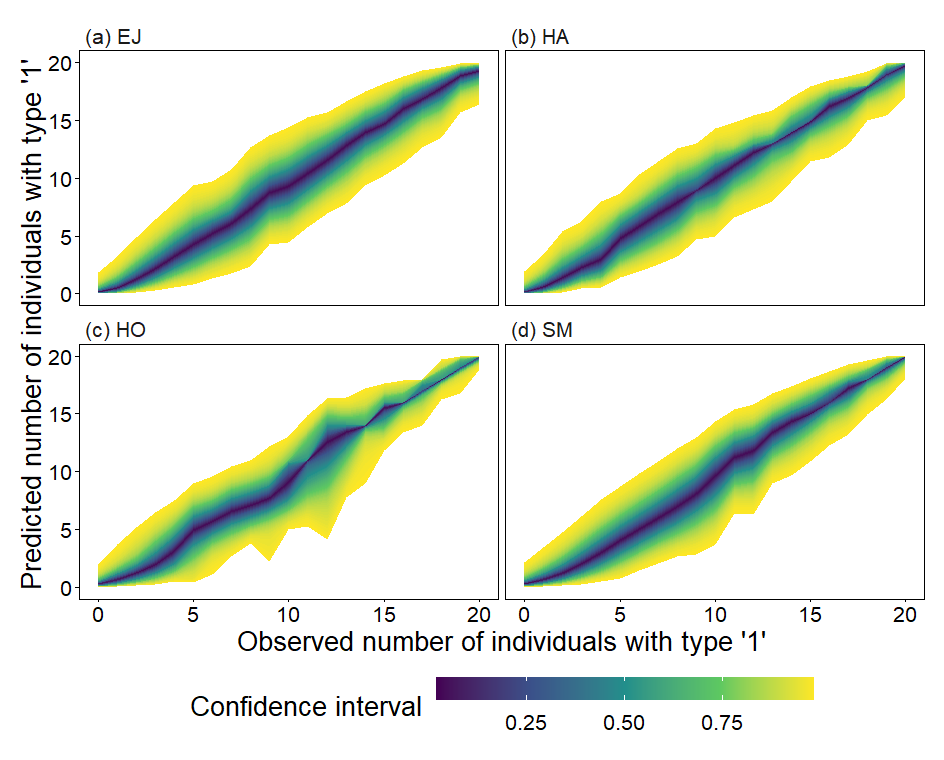

**Figure S4.** Predicted number of individuals with allelic type ‘1’ for an observed one at each locus from the total number of individuals (with types ‘1’ and ‘0’ together) in each local population (see Equation 1) for (a) *Ephemera japonica* (EJ), (b) *Hydropsyche albicephala* (HA), (c) *Hydropsyche orientalis* (HO) and (d) *Stenopsyche marmorata* (SM) with posterior distribution in Bayesian modelling. The detail of color scale is shown in the legend of Figure S1.

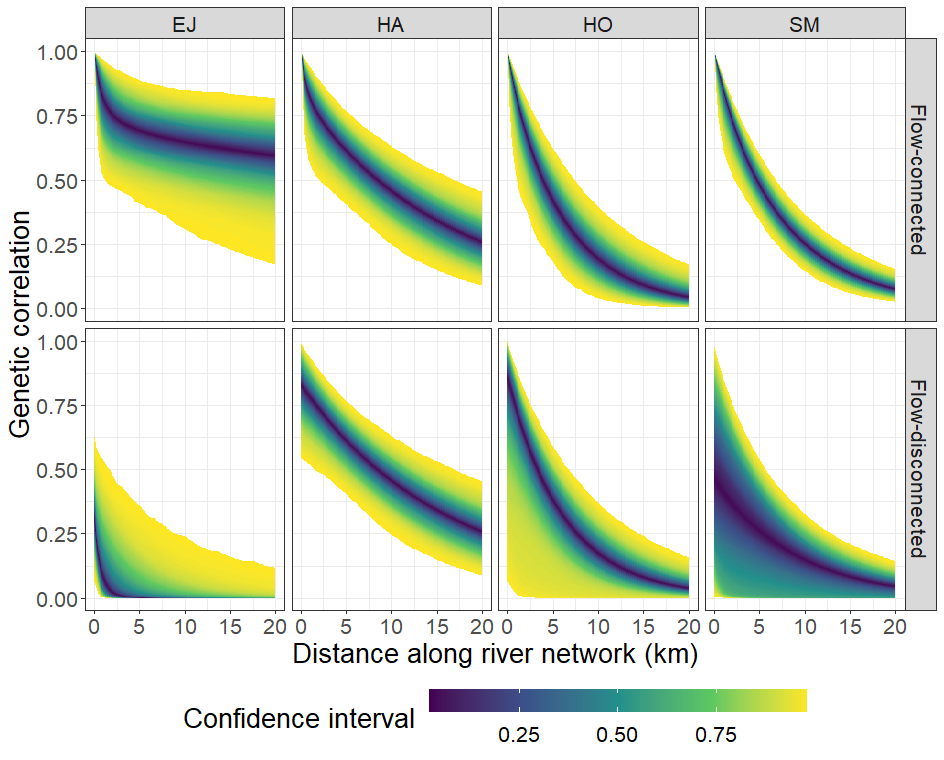

**Figure S5.** Change in genetic correlation by distance (covariance $\Omega_{ij}$ divided by variance $\sigma_{D}^{2}+\sigma_{U}^{2}$ in the genetic covariation function; with the posterior distribution in Bayesian modeling; see Equation 4) along river network between any two different streamflow-connected or -disconnected locations for *Ephemera japonica* (EJ), *Hydropsyche albicephala* (HA), *Hydropsyche orientalis* (HO) and *Stenopsyche marmorata* (SM). The detail of color scale is shown in the legend of Figure S1.

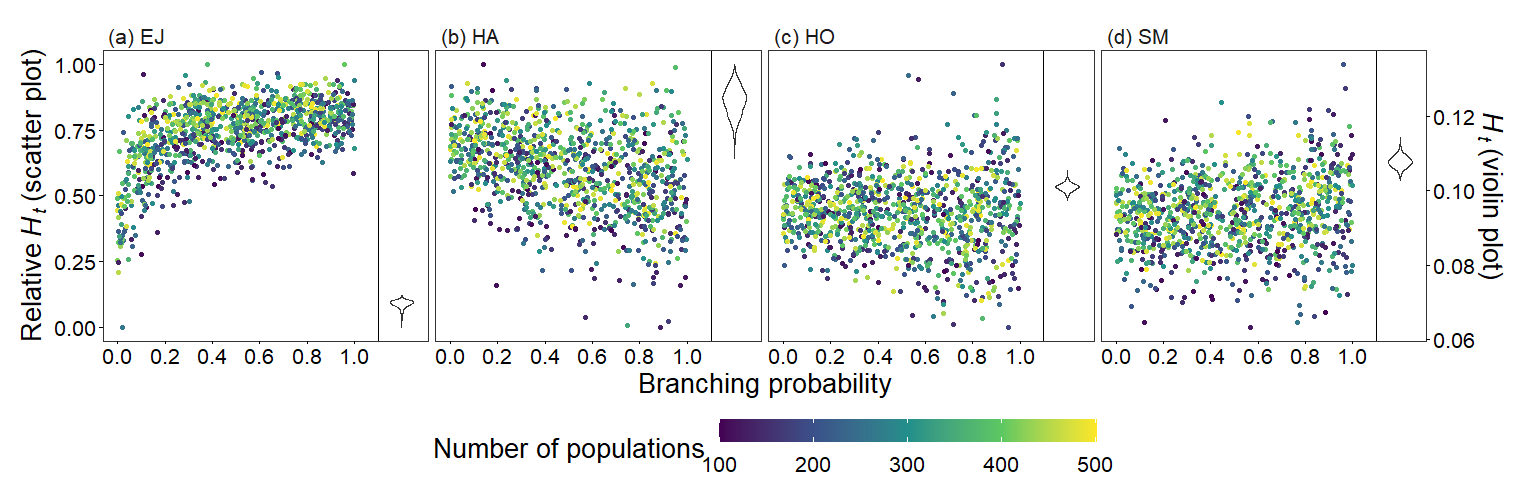

**Figure S6.** Theoretical predictions for relationships between genetic diversity (global $H_{t}$ = $GH_{t}$) and branching complexity under differential metapopulation sizes (range: 100 to 500, number of local populations) for (a) *Ephemera japonica* (EJ), (b) *Hydropsyche albicephala* (HA), (c) *Hydropsyche orientalis* (HO) and (d) *Stenopsyche marmorata* (SM). Relative ${GH}_{t}$ = ($GH_{t}$ – minimum $GH_{t}$) / (maximum $GH_{t}$ - minimum $GH_{t}$) across the range for each species.

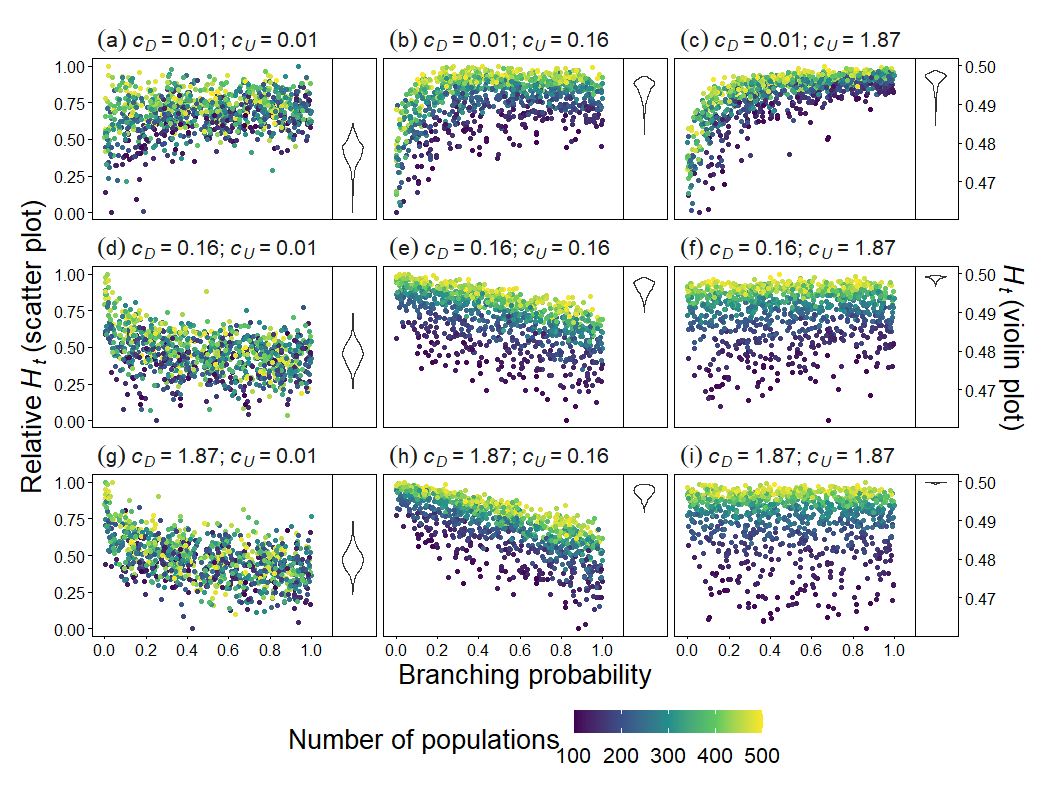

**Figure S7.** Theoretical predictions for relationships between metapopulation genetic diversity (global $H_{t}$ = $GH_{t}$) and branching complexity under differential metapopulation sizes (range: 100 to 500, number of local populations) for combinations of dispersal-related scale parameters in genetic covariation function (Equation 4), including (a, e and i) $c_{D}=c_{U}$, (b, c and f) $c_{D}<c_{U}$, and (d, g and h) $c_{D}>c_{U}$. The three values of each parameter are the upper, median, and lower ends of ranges of Bayesian-median pooled estimates across the four macroinvertebrate species. Relative ${GH}_{t}$ = ($GH_{t}$ – minimum $GH_{t}$) / (maximum $GH_{t}$ - minimum $GH_{t}$) across the range for each parameter combination.

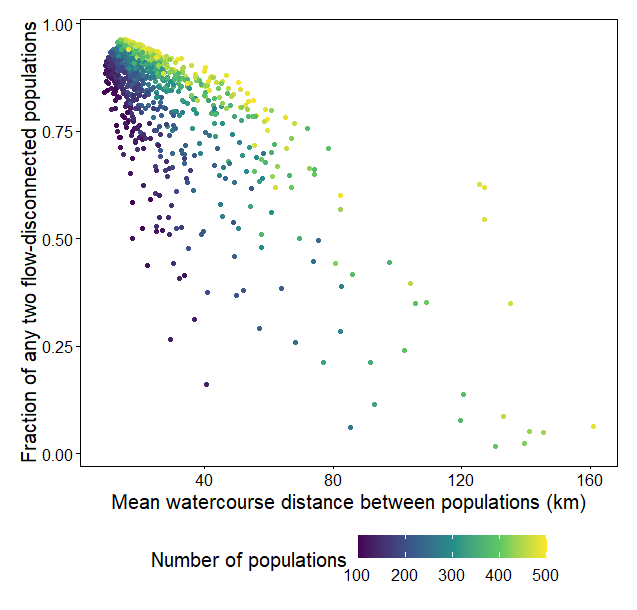

**Figure S8.** Theoretical predictions for relationships of the mean watercourse-distance between populations and the fraction of any two streamflow-disconnected populations (e.g., in headwaters) under differential metapopulation sizes (range: 100 to 500 local populations).

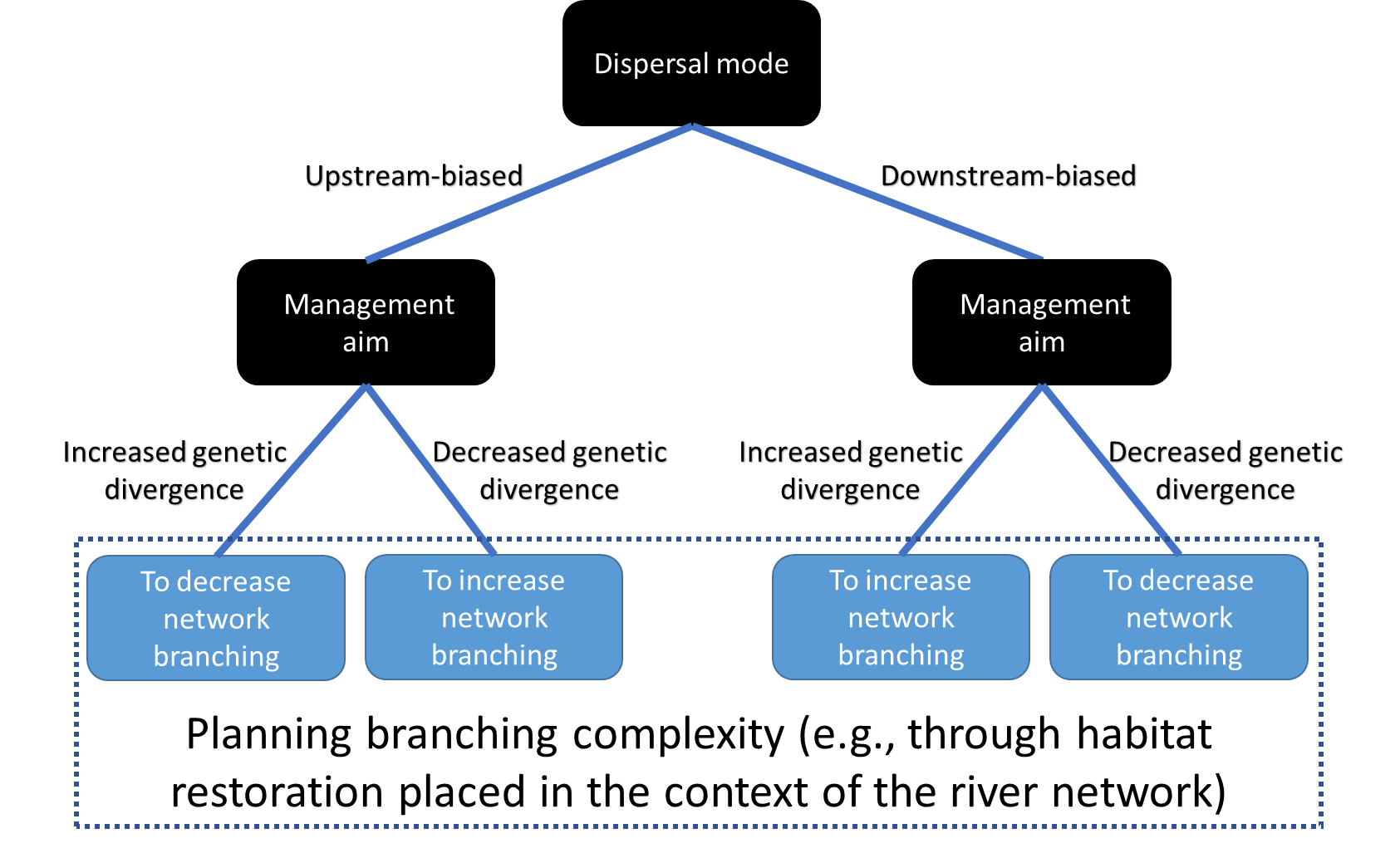

**Figure S9.** Decision tree for planning branching complexity based on species dispersal modes and management aims.
